# Fronto-Parietal Interactions with Task-Evoked Functional Connectivity During Cognitive Control

**DOI:** 10.1101/133611

**Authors:** Kai Hwang, James M. Shine, Mark D’Esposito

## Abstract

Flexible interaction between brain regions enables neural systems to transfer and process information adaptively for goal-directed behaviors. In the current study, we investigated neural substrates that interact with task-evoked functional connectivity during cognitive control. We conducted a human fMRI study where participants selectively attended to a category of visual stimuli in the presence of competing distractors from another stimulus category. To study flexible interactions between brain regions, we performed a dynamic functional connectivity analysis to estimate temporal changes in connectivity strength between brain regions under different levels of cognitive control. Consistent with theoretical predictions, we found that cognitive control selectively enhances functional connectivity for prioritizing the processing of task-relevant information. By regressing temporal changes in connectivity strength against activity patterns elsewhere in the brain, we localized frontal and parietal regions that potentially provide top-down biasing signals for influencing, or reading information out from, task-evoked functional connectivity. Our results suggest that in addition to modulating local activity, fronto-parietal regions could also exert top-down biasing signals to influence functional connectivity between distributed brain regions.

## Introduction

Cognitive control refers to our ability to flexibly regulate behavior to achieve changing goals. It is proposed that neural systems implement cognitive control by exerting top-down biasing signals to adaptively modulate on-going sensorimotor functions (Miller and Cohen, 2001). It is well established that top-down biasing signals can modulate the gain of localized brain activity (O’Craven et al., 1999; Treue and Trujillo, 1999), resulting in both enhancement of responses to task-relevant stimuli and dampening of responses to task-irrelevant stimuli (Gazzaley et al., 2005). These biasing signals can also modulate patterns of activity within local functional brain regions (Seidl et al., 2012; Nelissen et al., 2013). For example, goal-directed attention influences the decoding accuracy of multi-voxel patterns of activity in occipito-temporal cortices and its tuning to attended stimuli (Serences et al., 2009; Chen et al., 2012). These control mechanisms can adaptively change the signal strength and signal-to-noise ratio of information depending on its behavioral relevance, which is proposed to resolve the competition among stimuli or response pathways (Desimone, 1998; Badre, 2008). The prefrontal cortex (PFC) is a likely source of top-down biasing signals (Miller and D’Esposito, 2005). For example, patients with PFC lesions showed increased distractor-related evoked responses in posterior cortices (Knight et al., 1999), and temporary disruption of PFC function with transcranial magnetic stimulation (TMS) in healthy individuals reduces the gain of stimulus-evoked responses and decreases the discriminability of multivariate response patterns of activity in posterior cortices (Higo et al., 2011; Zanto et al., 2011; Lee and D’Esposito, 2012).

Neural systems not only support cognition through modulating localized information representation, but also through interactive communication between distributed brain regions (Van Essen et al., 1992; Friston, 2009). For example, while attending to a visual object, information related to elementary visual features encoded in primary visual cortex is transmitted to anterior ventral temporal cortices for further processing (Lerner et al., 2001). This suggests that cognitive control could also be achieved by adaptively regulating the information flow between brain regions to prioritize the transfer of task-relevant information (Botvinick et al., 2001; Miller and Cohen, 2001). However, few studies have investigated on how biasing signals influence information exchange between distributed brain regions.

Inter-regional communication between brain regions has been hypothesized to be a key mechanism for brain information processing (Bullmore and Sporns, 2009), and can be quantified by calculating the statistical dependency between activity in different brain regions (otherwise known as functional connectivity). Most methods that estimate functional connectivity from brain imaging data, such as Pearson correlation or coherence, assume that connectivity is stationary across time. However, recent studies have demonstrated that patterns of functional connectivity not only change across time (Hutchison et al., 2013), the temporal dynamics of functional connectivity are modulated by alterations in attention (Shine et al., 2016a) and memory (Braun et al., 2015) processes. In this human fMRI study, we used a dynamic functional connectivity method (Shine et al., 2015) to estimate, with high temporal resolution, patterns of task-evoked functional interactions between primary and higher-level visual areas during selective attention to categories of visual stimuli. As predicted by theoretical models and previous findings (Miller and Cohen, 2001; Al-Aidroos et al., 2012), we hypothesized that cognitive control should enhance information exchange between visual areas to prioritize task-relevant information processing. Further, by regressing the time point by time point dynamic functional connectivity estimates against whole brain activity, we were also able to identify localized regions that may putatively act as the source of top-down biasing signals that influence task-evoked functional connectivity patterns.

## Methods

### Subjects

Twenty-nine healthy adults subjects were recruited for this study. Four subjects were excluded due to excessive head motion, thus we report data from 25 subjects (aged 18-35, mean = 21.22, *SD* = 2.44, 15 males). All subjects were right handed, had normal or corrected to normal vision and reported no history of a neurological or psychiatric disorder. All patients provided written informed consent in accordance with procedures approved by the Committee for the Protection of Human Subjects at the University of California, Berkeley.

### Data Acquisition

Imaging data were acquired using a Siemens Tim/Trio 3T scanner and a 12-channel head coil. Structural images were acquired using a multi-echo MPRAGE sequence (TR = 2530ms; TE = 1.64/3.5/5.36/7.22 ms; flip angle = 7°; field of view = 256x 256, 176 sagittal slices, 1 mm^3^ voxels; 2x GRAPPA acceleration). Functional images were acquired using an echo-planar sequence sensitive to blood oxygenated level-dependent (BOLD) contrast (TR =1500 ms; TE = 25 ms; flip angel = 70°; field of view = 256x 256, voxel size: 4mm^3^ isotropic voxels with 29 contiguous axial slices in descending order; no acceleration). Each subject completed 24 runs of functional scans, each run lasting 2 minutes and 33 seconds (102 volumes each, total = 2448 volumes; total scan time approximately 75 minutes). An LCD projector projected visual stimuli onto a screen mounted to the head coil. Psychophysics Toolbox Version 3 was used to present stimuli and record responses and via a fiber-optic motor response-recording device.

### Experimental Task

Each subject completed 8 runs of a functional localizer task, and 16 runs of a two-factor experimental task (Figure 1). The localizer task was used to independently identify regions of interests (ROIs) for the connectivity analyses. We used pictures of human faces for localizing the fusiform face area (FFA; Kanwisher et al., 1997), pictures of buildings for the parahippocampal place area (PPA; Epstein et al., 1999), and both categories for localizing the primary visual cortex (V1). The two factors for the main experimental task were conditions (categorization task versus 1-back task) and the stimulus categories (faces versus buildings). In all tasks, subjects were asked to respond to sequences of pictures. In the localizer task and categorization task, subjects categorized pictures presented. For the 1-back task, subjects detected occasional repetitions of one stimulus category in the presence of competing stimuli from another category, thus requiring more cognitive control resources compared to the categorization task (Gazzaley et al., 2005; Lee and D’Esposito, 2012). The interaction between the 1-back task and stimuli categories (face versus building) further manipulated behavioral relevance of the stimuli. For example, a face was a relevant stimulus, and a building was an irrelevant stimulus in the 1-back attend to face condition.

**Figure 1.**
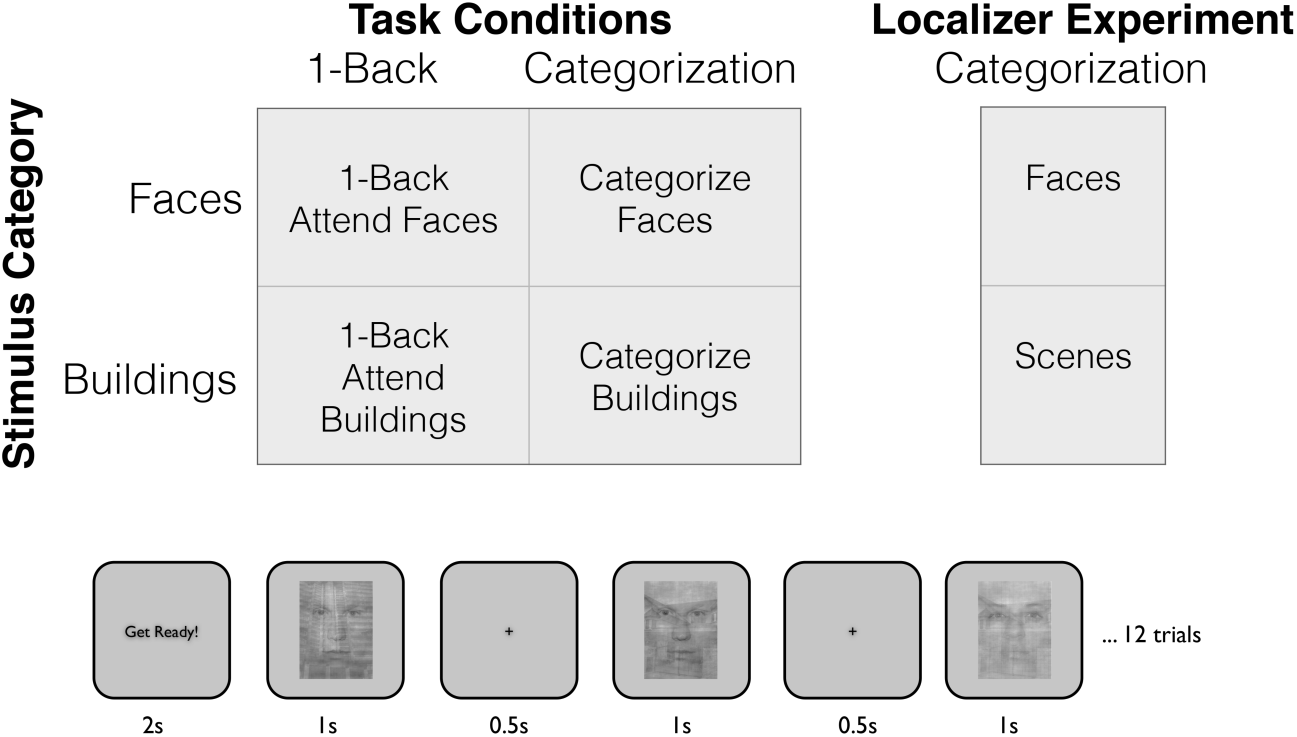
Structure of the task and trial timing.

All tasks began with 3 seconds of initial fixation, followed by 4 task blocks (20 seconds each) interleaved with 3 baseline fixation blocks (20 seconds each), and a 10 seconds fixation block (Figure 1). Each block started with a 2 seconds initiation cue. Task blocks consisted of 12 trials of stimuli. Each trial started with an image presented centrally on screen for 500 ms, immediately followed by a 1 second of fixation. For the localizer task, pictures of human faces or buildings were presented, and subjects were required to make a button press to identify the category of the picture (face or building). For the 1-back task, images presented were semi-transparent faces overlaid on semi-transparent phase-scrambled buildings, or semi-transparent buildings overlaid on semi-transparent phase-scrambled faces. Subjects were instructed to attend to a target category (faces or buildings), and were required to make a button press to the occasional back-to-back repetitions of this target category. There were 2 to 4 repetitions for both the attended and unattended categories within each block, and the presentation sequences were randomized separately. For the categorization task, images were semi-transparent faces or buildings overlaid on semi-transparent phase-scrambled faces or buildings from the opposite category. The phase-scramble procedure ensured that elementary visual properties of stimuli used were equivalent between the 1-back and categorization tasks. Subjects were required to make a button press to indicate the stimulus category. Luminance for all pictures was equalized using the SHINE toolbox (Willenbockel et al., 2010). For all tasks, subjects used their left or right index finger to make the button press, and the hand order was randomized across runs. A feedback indicating the accuracy of responses from the previous run was given at the end of each run.

### Data Analysis

*Preprocessing.* Imaging data were processed using AFNI and FSL (Cox, 1996; Smith et al., 2004). Functional images were slice-time and motion corrected (FSL’s slicetimer and MC-FLIRT), co-registered to T1-weighted structural image using a boundary-based-registration algorithm (FSL’s FLIRT), and warped to the MNI template using FSL’s non-linear registration (FSL’s FNIRT). Functional data were then resampled to 2mm combining motion correction and atlas transformation in a single interpolation step. Data were then spatially smoothed with a 5mm full-width-at-half-maximum Gaussian kernel (FSL’s SUSAN). We then performed nuisance regression using ordinary least squares regression (AFNI’s 3dDeconvolve) with the following regressors: polynomial fits for removing linear drifts, six rigid-body motion parameters and its derivatives, averaged signal from white-matter and ventricles ROIs created using freesurfer’s tissue segmentation tool (Dale et al., 1999). To minimize motion confounds, we calculated average frame-wise displacement (FD) (Power et al., 2012), and volumes with FD > 0.2 were excluded from all analyses. Subjects with > 25% volumes removed were also excluded (n=4).

*Regions of interets (ROIs) analyses.* After preprocessing, we performed a generalized linear model (GLM) analysis of linear regression at each voxel, using generalized least squares with a voxel-wise ARMA(1,1) autocorrelation model (AFNI’s 3dREMLfit). Finite impulse response (FIR) basis functions were used to estimate the mean stimulus-evoked response amplitudes during task blocks, separately for each condition (localizers, categorize and 1-back conditions crossed with stimulus categories). Using data from the localizer runs, we then defined the FFA/PPA using the top 255 voxels (size equivalent to a 8 mm sphere) that were most selective for faces (face blocks > building blocks) and buildings (building blocks > faces blocks) within previously defined FFA/PPA ROIs (Julian et al., 2012). We defined the V1 ROI using the top 255 most active voxels within an anatomically defined ROI of each individual subject’s calcarine sulcus (Destrieux et al., 2010). To analyze differences in stimulus-evoked response amplitudes, for each ROI we performed a three-way within subject analysis of variance, crossing conditions (1-back vs. categorize), stimulus categories (faces vs. buildings) and time (21 volumes within each task block).

*Functional connectivity analyses.* Our goal was to localize potential sources of top-down biasing signals that modulate task-evoked functional connectivity patterns. Achieving this goal requires a method that can estimate temporal changes in connectivity patterns under different task conditions, which can then be used to localized regional changes in brain activity that covary with temporal changes in connectivity estimates. Such a method complements existing static functional connectivity methods that assume connectivity structure is stationary and invariant across time and task conditions. Thus, we utilized a dynamic functional connectivity metric, Multiplication of Temporal Derivatives (MTD; for details see Shine et al., 2015), to estimate time varying connectivity strength between V1 and PPA/FFA under task conditions. Prior to preforming all connectivity analyses, stimulus-evoked responses were regressed out from the preprocessed data, and residuals were used to assess task-evoked functional connectivity independent of shared variances between ROIs (Norman-Haignere et al., 2012; Cole et al., 2014; Gratton et al., 2016). This additional regression was performed to minimize the influence of mean task-related activation on task-evoked functional connectivity, while retaining the residual trial-by-trial fluctuations in the time-series that contributes to task-evoked functional connectivity. To perform MTD analysis, we first calculated the first order temporal derivatives (dt) of each time-series extracted from ROIs, and then normalized each data point by dividing each derivative by the standard deviation of the whole time-series. We then multiplied dt scores to calculate MTD scores between ROIs. Positive MTD scores reflect synchronized coupling between ROIs, whereas negative MTD scores reflect out-of-synch decoupling. Similar to a Pearson correlation analysis, the MTD values can be averaged across time and be interpreted as static functional connectivity strength, whereas the time-varying MTD scores reflect dynamic changes in functional connectivity.

Because MTD scores of a single time point could be susceptible to high-frequency noise, we further calculated a simple moving average on MTD scores (Shine et al., 2015). To determine the most effective window length for detecting task-evoked functional connectivity (i.e. connectivity between time-series that occurred primarily within task-blocks), we simulated time-series (Figure 2A) that contained: (i) 1^st^ order correlations over the entire task (task and resting blocks); and (ii) 2^nd^ order correlations that occurred only during the task blocks (i.e. task correlations). We ran 5000 iterations of this simulation and calculated the MTD across a range of window lengths from 10 volumes to 100 volumes. We then determined the effect size of the difference between the task-evoked and resting correlation time-series (Figure 2B). Windows with the length of 15 volumes showed the maximal effect size for differentiating 1^st^ order and 2^nd^ order functional connectivity. Therefore all results were presented using a smoothing window of 15 volumes, but we also explored a range of smoothing windows (1 to 20). Importantly, MTD time-series for each condition can be used as additional regressors in the GLM analysis to localize potential sources of signals that modulate task-evoked functional connectivity patterns.

**Figure 2.**
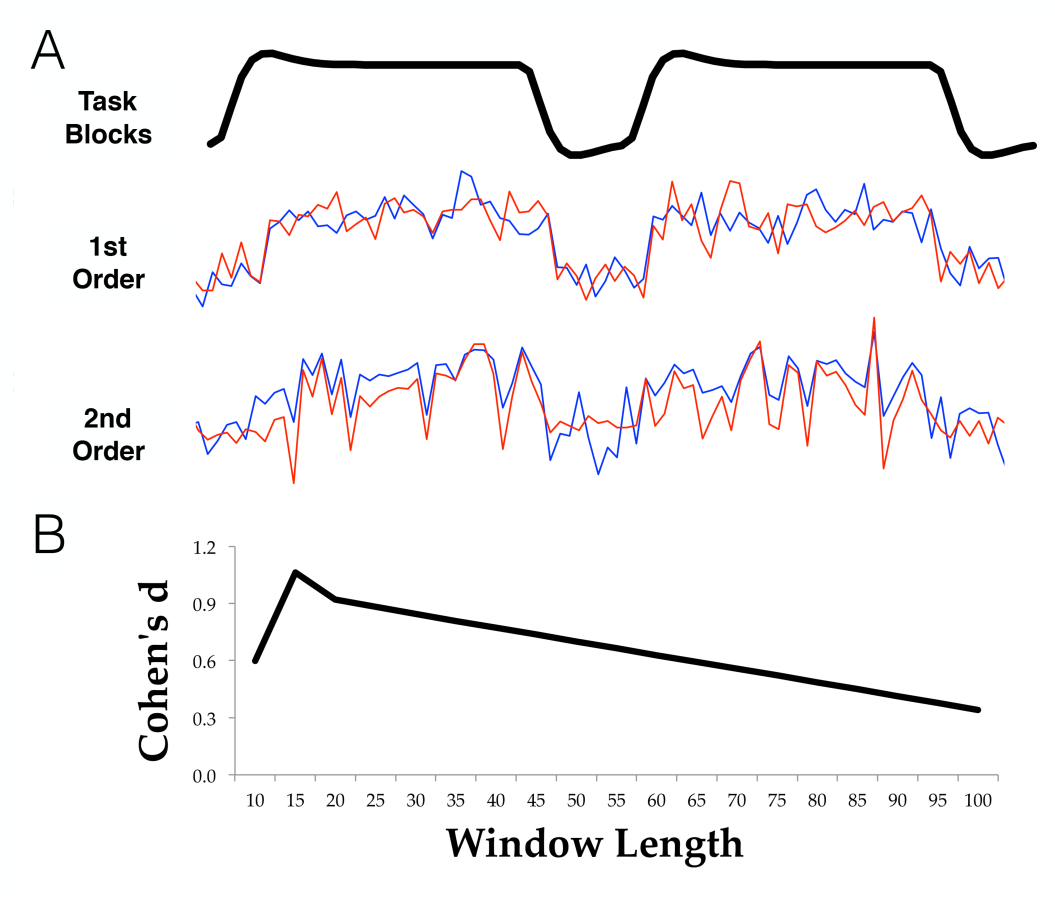
Simulation for determining optimal smoothing window. (A) Simulated time-series that have r = 0.8 correlations for both task and resting blocks (1^st^ order) or only during task blocks (2^nd^ order). (B) Effect size of differentiating between 1^st^ and 2^nd^ order correlations.

Whole-brain GLM maps of each individual subject’s MTD regressors were then submitted to group analysis contrasting the effects of condition (1-back versus categorize) and its interaction with stimulus relevance (relevant stimuli, e.g., V1-FFA connectivity during attend to face condition, versus irrelevant stimuli, e.g., V1-FFA connectivity during attend to building condition). Group level analysis was performed with a linear mixed effects model at each voxel, using generalized least squares with a local estimate of random effects variance (AFNI’s 3dMEMA). To correct for multiple comparisons, a Monte Carlo simulation of 5000 iterations was performed to identify minimal cluster size that reached a corrected family-wise error rate of 0.05 (AFNI’s 3dClustSim). The smoothness that entered this simulation was estimated from the GLM model residuals, using a spatial autocorrelation function (Cox et al., 2016) implemented in AFNI’s 3dFWHMx (full-width-half-maximum smoothness = 8.32 mm). Cluster forming threshold were set as *p* < 0.05, and the resulting minimal cluster size was 479 contiguous 2mm^3^ voxels. All unthresholded statistic maps have been uploaded to the NeuroVault database (http://neurovault.org/collections/2474/). We also performed seed-based connectivity analyses by including the mean signals extracted from FFA and PPA as additional regressors in the GLM model. The associated maps were submitted to group analysis using the same procedure.

*Connectivity and stimulus-evoked responses.* We tested the relationship between MTD estimates and stimulus-evoked response amplitudes by calculating its correlations across subjects. The amplitudes of each condition’s stimulus-evoked responses were calculated using area-under-the-curve of FIR estimates. We further correlated subject-by-subject seed-based connectivity strength estimates (Pearson correlations with FFA/PPA) from selected fronto-parietal ROIs with stimulus-evoked response amplitudes. These ROIs were created by creating a 5mm radius sphere centered on peak voxels in each significant spatial cluster that showed significant connectivity with FFA/PPA (right inferior precentral sulcus: x = 20, y = −8, z = 56; right insula: x = 42, y = 24, z = 0; right intraparietal sulcus: x = 26, y = −58, z = 60; left middle frontal: x = 18, y = −36, z = 50).

## Results

### Behavioral Results

Behavioral performance for the categorization task was near ceiling for both faces and buildings (mean accuracy for faces = 0.96, *SD* = 0.08; mean accuracy for buildings = 0.96, *SD* = 0.06; mean RT for faces = 543 ms, *SD* = 99 ms; mean RT for buildings = 557 ms, *SD* = 109 ms). Accuracy for detecting repetitions in the 1-back condition was below ceiling, and no significant difference was found between faces and buildings (mean accuracy for faces = 0.71, *SD* = 0.16; mean accuracy for buildings = 0.74, *SD* = 0.16; mean RT for faces = 745 ms, *SD* = 112 ms; mean RT for buildings = 693 ms, *SD* = 100 ms; *t*-test for accuracy: *t*(24) = 1.07, *p* = 0.29; *t*-test for RT: *t*(24) = 7.39, *p* = 1.43×10^-7^). The accuracy for 1-back (only including repeating stimuli) was significantly lower when compared to the categorize task (faces: *t*(24) = 6.71, *p* = 7.27×10^-7^; buildings: *t*(24) = 1.02, *p* =1.02×10^-7^). Similarly, reaction time for 1-back task was significantly slower when compared to the categorization task (faces: *t*(24) = 13.59, *p* = 7.17×10^-13^; buildings: *t*(24) = 8.39, *p* = 1.45×10^-8^).

### Stimulus Evoked Responses

We found increased stimulus-evoked responses in V1 for all task conditions (Figure 3A, main effect of factor volume: *F*(1,2084) = 9.38, *p* = 0.0022), but no significant condition by category interaction (*F*(1,2084) = 0.31, *p* = 0.56). We found significant condition by category interactions in both the FFA and PPA (Figure 3B-C; FFA: *F*(1,2084) = 4.21, *p* = 0.04; PPA: *F*(1,2084) = 4.87, *p* = 0.027). Specifically, averaged across time points within task blocks, faces elicited stronger stimulus-evoked responses in FFA compared to buildings (t(24) = 11.71, *p* = 1.82×10^-11^), whereas buildings elicited a stronger responses in PPA compared to faces (t(24) = 11.61, *p* = 2.18×10^-11^). Further, for both the FFA and PPA, the 1-back condition elicited stronger stimulus-evoked responses compared to the categorize condition (FFA: *t*(24) =4.29, *p* = 0.00032; PPA: *t*(24) = 5.39, *p* = 1.95×10^-5^).

**Figure 3.**
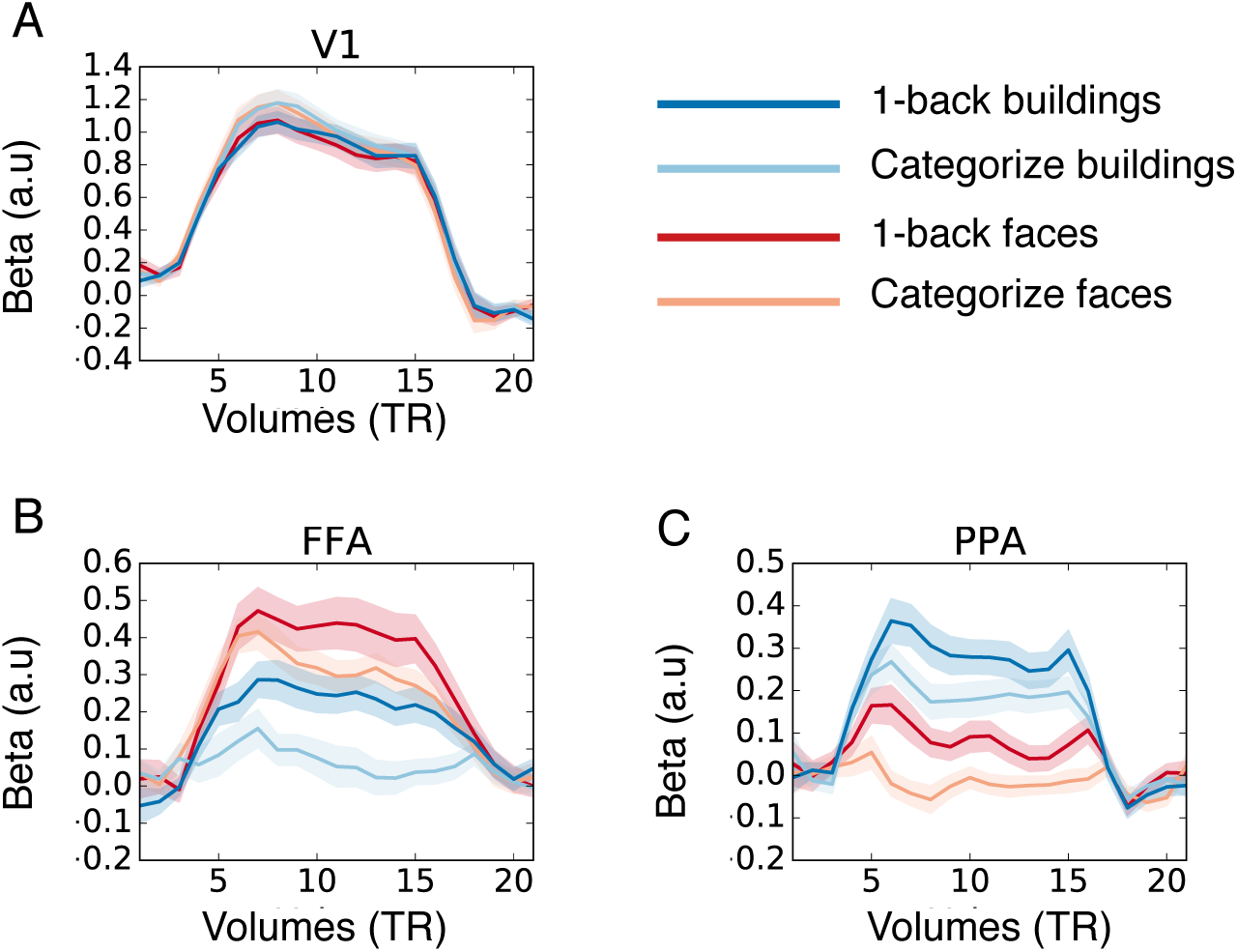
Stimulus-evoked responses. Y axis indicates stimulus-evoked response magnitude, X axis indicates the duration of task blocks. Shaded areas represent 1 SE.

### Dynamic Functional Connectivity

Based on theoretical models and previous findings (Cohen and Miller, 2001; Al-Aidroos et al., 2012), we predicted that functional connectivity between V1 and ventral visual cortex should be category-specific. That is, when subjects attend to faces, functional connectivity between V1 and FFA should increase compared to when they attended to buildings. In contrast, attention to buildings should increase connectivity between V1 and PPA when compared to attention to faces. Consistent with our predictions, we found that functional connectivity (calculated using time-averaged MTD scores) between task-related brain regions is modulated by cognitive control and stimuli relevance (Figure 4A). Specifically, functional connectivity between V1 and PPA was stronger for the 1-back attend to buildings condition compared to the 1-back attend to faces condition (*t*(24) = 6.51, *p* = 1.18×10^-6^) and categorization conditions (1-back buildings versus categorize faces: *t*(24) = 5.98, *p* = 4.4×10^-6^; 1-back buildings versus categorize buildings: *t*(24) = 3.27, *p* = 0.0032). Functional connectivity between V1 and FFA was significantly stronger for the 1-back attend to faces condition when compared to the 1-back attend to buildings condition (*t*(24) = 4.44, *p* = 0.00017) and categorize building condition (1-back buildings versus categorize faces: *t*(24) = 1.98, *p* = 0.059; 1-back buildings versus categorize buildings: *t*(24) = 3.10, *p* = 0.003). Connectivity between V1 and FFA was stronger than V1 and PPA under the 1-back attend to faces condition (*t*(24) = 4.17, p = 0.00034). No significant difference was found for connectivity between V1-PPA and V1-FFA under the 1-back attend to buildings condition (*t*(24) = 0.17, *p* = 0.86). These results were consistent across a range of smoothing windows we explored (Figure 4B), and further consistent with connectivity strength estimated using Pearson correlations (Figure 4C). Furthermore, we did not find any significant correlations between evoked-response amplitudes and MTD estimates. MTD estimates for 1-back conditions were moderately correlated with subject’s accuracy on the 1-back task (1-back attend to face: *r*(24) = 0.37, *p* = 0.075; 1-back attend to buildings *r*(24) = 0.44, *p* = 0.031).

**Figure 4.**
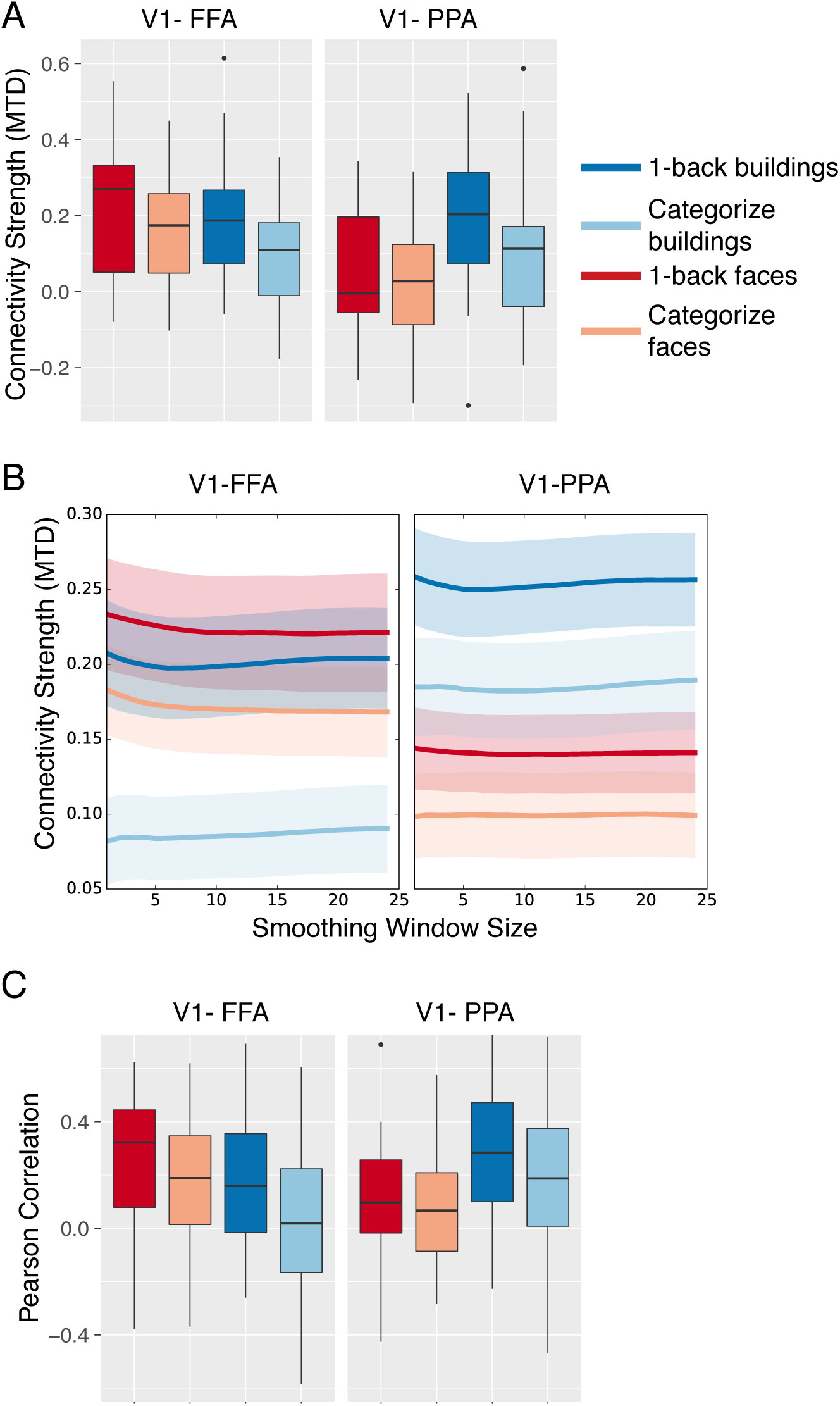
Task-evoked functional connectivity. (A) Box-plots of functional connectivity strength (MTD scores) between V1 and FFA/PPA under different experimental conditions. Box plot percentiles (5^th^ and 95^th^ for outer whiskers, 25^th^ and 75^th^ for box edges, and median for horizontal line with the median) were calculated across subjects separately for each condition. Single dots represent values outside of 5^th^ and 95^th^ percentiles. (B) MTD scores across a range of smoothing window sizes. Thick solid lines represent the mean MTD values of each condition. Shaded areas represent 1 SE. (C) Box plots of Pearson correlations calculated between V1/FFA for different conditions.

To localize potential sources of top-down biasing signals for modulating task-evoked functional connectivity, we entered each condition’s time-varying MTD scores as additional regressors in the whole-brain GLM analysis and contrasted 1-back versus categorization conditions. We found that a distributed set of fronto-parietal regions that showed increased activity that was positively associated with changes in task-evoked connectivity for processing task-relevant stimuli (i.e., averaging MTD estimates of V1-FFA for attend faces condition and V1-PPA for attend buildings conditions; Figure 5A). These regions included bilateral superior frontal sulcus, the left middle frontal gyrus, bilateral dorsal medial frontal cortex, bilateral precuneus, and bilateral intraparietal sulcus. Negative associations were found in the medial occipital cortex. This indicates that increased activity in these frontal and parietal regions is associated with increases in connectivity strength between V1 and higher order visual areas for processing the attended visual stimuli. No significant clusters of activation associated with processing of task-irrelevant stimuli were found after correcting form multiple comparisons.

**Figure 5.**
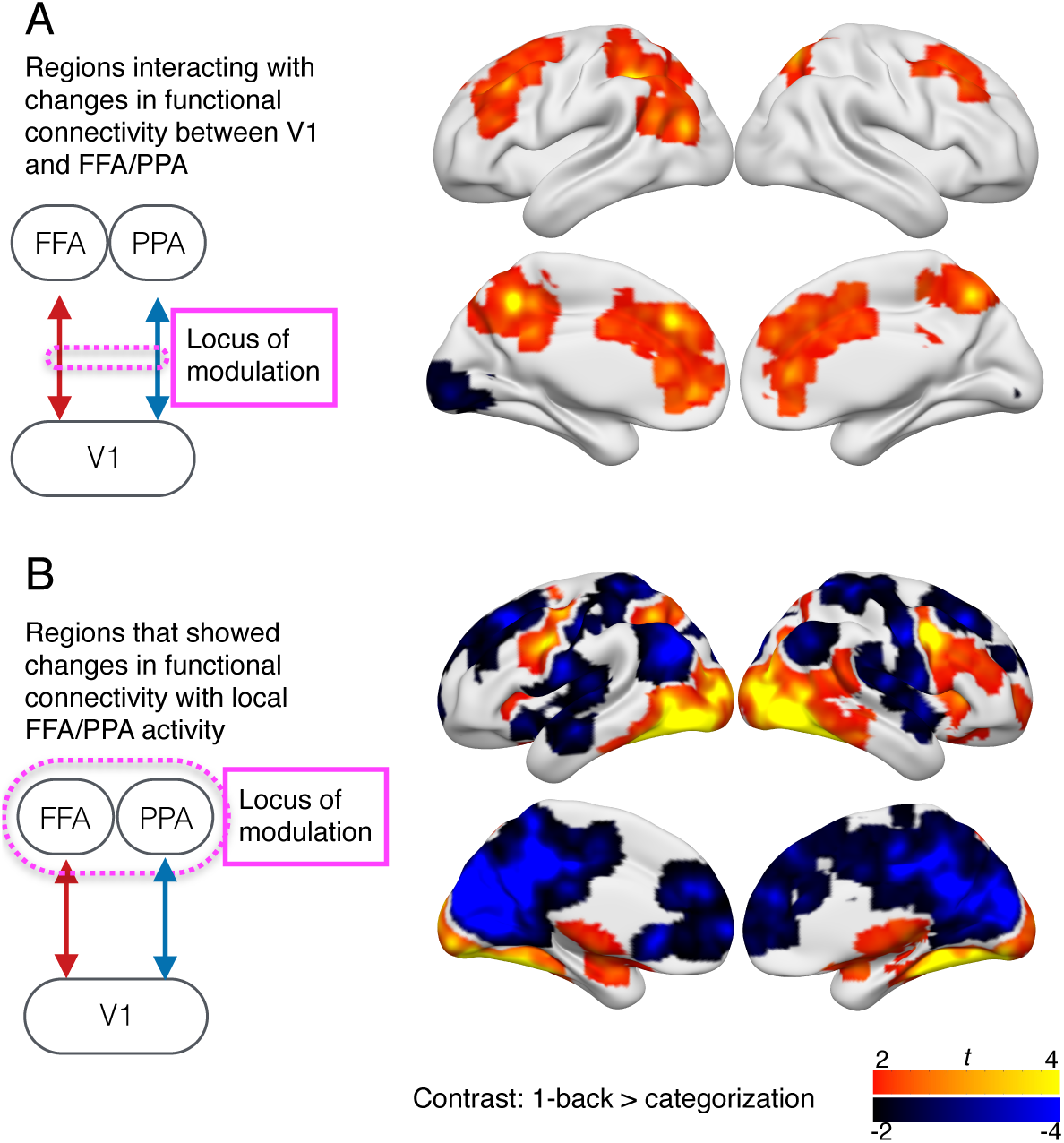
Potential sources of top-down biasing signals. (A) Regions that showed significant task modulations in interactions with task-evoked connectivity patterns between V1 and FFA/PPA. This map was generated using a smoothing window of 15 volumes. (B) Regions that showed significant task-related changes in functional connectivity with local FFA/PPA activity.

### Seed-Based Functional Connectivity

It is possible that the fronto-parietal brain regions identified in the previous analysis were simultaneously interacting with FFA/PPA and V1, but not specifically modulating functional connectivity strength between FFA/PPA and V1. To rule out this possibility, we performed a seed-based functional connectivity analysis to localize brain regions that showed task-driven changes in connectivity (contrasting 1-back versus categorize conditions) with FFA/PPA. In this analysis, we found increased connectivity with bilateral inferior precentral sulcus, the right inferior frontal gyrus, bilateral insular, the left intraparietal sulcus, bilateral occpito-temporal cortices, and the thalamus. We further found decreased connectivity with bilateral superior frontal sulcus, bilateral central sulcus, the dorsal medial prefrontal cortex, bilateral inferior parietal cortex, bilateral medial occipital cortex, bilateral superior temporal cortex, and bilateral medial parietal cortex (Figure 5B). Importantly, except the intraparietal sulcus, most brain regions that showed increased functional connectivity with FFA/PPA (Figure 5A) exhibited little overlap with brain regions that showed a positive association with dynamic functional connectivity patterns (Figure 5B). Instead, brain regions that showed decreased functional connectivity with FFA/PPA overlapped with brain regions that showed a positive association with dynamic functional connectivity patterns. No significant correlations were found between seed-based connectivity estimates and behavioral performance.

## Discussion

In this study, we investigated the neural mechanisms underlying top-down biasing signals that influence task-evoked functional connectivity. Utilizing a dynamic functional connectivity approach that allowed us to localize regions that co-vary with temporal fluctuations in functional connectivity, we found that task-related changes in functional connectivity patterns interact with activity in several frontal and parietal regions. Our results suggest that fronto-parietal regions could be the source of top-down signals that bias information communication along functional pathways for cognitive control.

Previous studies of patient with focal lesions or healthy individuals following TMS have provided ample evidence that frontal and parietal cortices provide top-down biasing signals that influence activity in posterior sensory cortices (i.e., Armstrong and Moore, 2007; Ruff et al., 2008; Feredoes et al., 2011; Higo et al., 2011; Zanto et al., 2011; Lee and D’Esposito, 2012; Gregoriou et al., 2014; Heinen et al., 2014; Lorenc et al., 2015). For example, TMS of dorsal lateral prefrontal cortex (Feredoes et al., 2011), inferior frontal cortex (Lee and D’Esposito, 2012, Zanto et al., 2011), frontal eye fields (Heinen et al., 2014) and intraparietal sulcus (Ruff et al., 2008) can modulate occipito-temporal activity. In addition, stimulating the frontal cortex modulates the discriminability of neural population code in occipito-temporal cortices (Armstrong and Moore, 2007; Lee and D’Esposito, 2012; Lorenc et al., 2015). Likewise, lateral PFC lesions reduce the attentional effect on stimulus-evoked response amplitudes and neural synchrony in V4 (Gregoriou et al., 2014). In aggregate, these studies suggest that one major function of fronto-parietal cortices is to modulate the gain and tuning in posterior cortices in the service of cognitive control. These empirical findings support the theory of biased-competition (Desimone and Duncan, 1995), which proposes that selection of goal-relevant information in specialized brain regions are enhanced by top-down biasing signals.

Our results demonstrated that another potential function of fronto-parietal cortices is to adaptively modulate functional connectivity between brain regions, putatively to establish signal pathways for transferring task-relevant information. There are at least two potential mechanisms that could support this function. First, top-down biasing signals emanating from fronto-parietal cortices could modulate neuronal oscillations between brain regions, which are proposed to facilitate communication via temporal coherence (Fries, 2015). It has been suggested that different classes of neural oscillations (i.e., alpha-, beta-, gamma-band oscillations) could be generated by distinct neurophysiological mechanisms (Moore et al., 2010; Wang, 2010). Neural oscillations are putatively controlled by neocortical inhibitory interneurons (Cardin et al., 2009; Vierling-Claassen et al., 2010), suggesting that top-down modulations of neural synchrony between brain regions may target cortical inhibitory neurons, thereby affecting brain region’s excitability to synaptic inputs and its communication efficacy (Lakatos et al., 2005). A second potential mechanism could involve fronto-parietal regions acting on the thalamus or the thalamic reticular nucleus (Wimmer et al., 2015), which in turn could modulate cortico-cortical communication. In addition to relaying information from peripheral sensory organs to the cerebral cortex, the thalamus has also been proposed to mediate the exchange of information between cortical regions through cortico-thalamic-cortical pathways (Sherman, 2016). For example, the pulvinar nucleus has been found to synchronize with distant visual areas according to attentional demands (Saalmann et al., 2012). These results suggest that the thalamus could receive top-down biasing signals from fronto-parietal cortices, and in turn be involved in regulating information communication between cortical regions according to various cognitive control demands.

Although the PFC has been proposed to support cognitive control by establishing the optimal route of information transfer between brain regions (Miller and Cohen, 2001), direct evidence has been lacking. Due to general assumptions of stationary connectivity over the course of an entire behavioral session, most existing studies have not examined contextual or temporal changes in functional connectivity under different cognitive control demands. New connectivity methods such as the one utilized here allowed us to provide such evidence (Al-Aidroos et al., 2012; Shine et al., 2015; Shine et al., 2016b). Given that we removed the mean trial-evoked responses in our analyses, our findings are not likely driven by correlated evoked response patterns. However, because the method we utilized is correlational, we cannot determine the direction of interaction between fronto-parietal regions and task-evoked functional connectivity in the visual cortex. It is possible that the fronto-parietal activity we observed reflects a read out mechanism for selecting goal-relevant information, rather than a top-down biasing signal. To address this possibility, we correlated signals extracted from FFA/PPA, and observed that fronto-parietal regions that exhibited increased static functional connectivity with FFA/PPA (Figure 5B) were different than the fronto-parietal regions that correlated with dynamic functional connectivity patterns between FFA/PPA and V1 (Figure 5A). We also found that estimates of dynamic functional connectivity patterns did not correlate with evoked response amplitudes, suggesting that mechanisms involved in local gain modulation could be dissociable from those from modulating task-evoked connectivity patterns between brain regions. Given previous work demonstrating fronto-parietal regions as the source of top-down biasing signals that modulate gain and tuning of activity in posterior cortical regions, we predict that these regions also provide a signal that can modulating task-adaptive connectivity. Future TMS or lesion studies can directly test this prediction.

In summary, our results suggest that there is likely a fronto-parietal mechanism for modulating the synchronizability between brain regions, which compliments other mechanisms such as bias competition (Desimone and Duncan, 1995), noise correlation (Cohen and Maunsell, 2009), and tuning change (Serences et al., 2009). Altogether, these mechanisms provide multiple means for brains to generate flexible and adaptive behaviors.

## References

Al-Aidroos N, Said CP, Turk-Browne >NB (2012) Top-down attention switches coupling between low-level and high-level areas of human visual cortex. Proc Natl Acad Sci U S A 109:14675–14680.

Armstrong KM, Moore T (2007) Rapid enhancement of visual cortical response discriminability by microstimulation of the frontal eye field. Proc Natl Acad Sci U S A 104:9499–9504.

Badre D (2008) Cognitive control, hierarchy, and the rostro-caudal organization of the frontal lobes. Trends Cogn Sci 12:193–200.

Botvinick MM, Braver TS, Barch DM, Carter CS, Cohen JD (2001) Conflict monitoring and cognitive control. Psychol Rev 108:624–652.

Braun U, Schafer A, Walter H, Erk S, Romanczuk-Seiferth N, Haddad L, Schweiger JI, Grimm O, Heinz A, Tost H, Meyer-Lindenberg A, Bassett DS (2015) Dynamic reconfiguration of frontal brain networks during executive cognition in humans. Proc Natl Acad Sci U S A 112:11678–11683.

Bullmore E, Sporns O (2009) Complex brain networks: Graph theoretical analysis of structural and functional systems. Nature Reviews Neuroscience 10:186–198.

Cardin J, Carlén M, Meletis K, Knoblich U, Zhang F, Deisseroth K, Tsai L, Moore C (2009) Driving fast-spiking cells induces gamma rhythm and controls sensory responses. Nature.

Chen AJ, Britton M, Turner GR, Vytlacil J, Thompson TW, D’Esposito M (2012) Goal-directed attention alters the tuning of object-based representations in extrastriate cortex. Front Hum Neurosci 6:187.

Cohen MR, Maunsell JH (2009) Attention improves performance primarily by reducing interneuronal correlations. Nat Neurosci 12:1594–1600.

Cole MW, Bassett DS, Power JD, Braver TS, Petersen SE (2014) Intrinsic and task-evoked network architectures of the human brain. Neuron 83:238–251.

Cox RW (1996) Afni: Software for analysis and visualization of functional magnetic resonance neuroimages. Comput Biomed Res 29:162–173.

Cox RW, Reynolds RC, Taylor PA (2016) AFNI and clustering: false positive rates redux. bioRxiv:065862.

Dale AM, Fischl B, Sereno MI (1999) Cortical surface-based analysis. I. Segmentation and surface reconstruction. NeuroImage 9:179–194.

Desimone R (1998) Visual attention mediated by biased competition in extrastriate visual cortex. Philos Trans R Soc Lond B Biol Sci 353:1245–1255.

Desimone R, Duncan J (1995) Neural mechanisms of selective visual attention. Annual Review of Neuroscience 18:193–222.

Destrieux C, Fischl B, Dale A, Halgren E (2010) Automatic parcellation of human cortical gyri and sulci using standard anatomical nomenclature. NeuroImage 53:1–15.

Epstein R, Harris A, Stanley D, Kanwisher N (1999) The parahippocampal place area: Recognition, navigation, or encoding? Neuron 23:115–125.

Feredoes E, Heinen K, Weiskopf N, Ruff C, Driver J (2011) Causal evidence for frontal involvement in memory target maintenance by posterior brain areas during distracter interference of visual working memory. Proc Natl Acad Sci U S A 108:17510–17515.

Fries P (2015) Rhythms for cognition: Communication through coherence. Neuron 88:220–235.

Friston KJ (2009) Modalities, modes, and models in functional neuroimaging. Science 326:399–403.

Gazzaley A, Cooney JW, McEvoy K, Knight RT, D’Esposito M (2005) Top-down enhancement and suppression of the magnitude and speed of neural activity. Journal of Cognitive Neuroscience 17:507–517.

Gratton C, Laumann TO, Gordon EM, Adeyemo B, Petersen SE (2016) Evidence for Two Independent Factors that Modify Brain Networks to Meet Task Goals. Cell Rep 17:1276–1288.

Gregoriou GG, Rossi AF, Ungerleider LG, Desimone R (2014) Lesions of prefrontal cortex reduce attentional modulation of neuronal responses and synchrony in V4. Nat Neurosci 17:1003–1011.

Heinen K, Feredoes E, Weiskopf N, Ruff CC, Driver J (2014) Direct evidence for attention-dependent influences of the frontal eye-fields on feature-responsive visual cortex. Cereb Cortex 24:2815–2821.

Higo T, Mars RB, Boorman ED, Buch ER, Rushworth MF (2011) Distributed and causal influence of frontal operculum in task control. Proc Natl Acad Sci U S A 108:4230–4235.

Hutchison RM, Womelsdorf T, Allen EA, Bandettini PA, Calhoun VD, Corbetta M, Della Penna S, Duyn JH, Glover GH, Gonzalez-Castillo J, Handwerker DA, Keilholz S, Kiviniemi V, Leopold DA, de Pasquale F, Sporns O, Walter M, Chang C (2013) Dynamic functional connectivity: promise, issues, and interpretations. Neuroimage 80:360–378.

Julian JB, Fedorenko E, Webster J, Kanwisher N (2012) An algorithmic method for functionally defining regions of interest in the ventral visual pathway. Neuroimage 60:2357–2364.

Kanwisher N, McDermott J, Chun MM (1997) The fusiform face area: A module in human extrastriate cortex specialized for face perception. J Neurosci 17:4302–4311.

Knight RT, Staines WR, Swick D, Chao LL (1999) Prefrontal cortex regulates inhibition and excitation in distributed neural networks. Acta Psychol (Amst) 101:159–178.

Lakatos P, Shah AS, Knuth KH, Ulbert I, Karmos G, Schroeder CE (2005) An oscillatory hierarchy controlling neuronal excitability and stimulus processing in the auditory cortex. J Neurophysiol 94:1904–1911.

Lee TG, D’Esposito M (2012) The dynamic nature of top-down signals originating from prefrontal cortex: A combined fmri-tms study. J Neurosci 32:15458–15466.

Lerner Y, Hendler T, Ben-Bashat D, Harel M, Malach R (2001) A hierarchical axis of object processing stages in the human visual cortex. Cereb Cortex 11:287–297.

Lorenc ES, Lee TG, Chen AJ, D’Esposito M (2015) The Effect of Disruption of Prefrontal Cortical Function with Transcranial Magnetic Stimulation on Visual Working Memory. Frontiers in systems neuroscience 9:169.

Miller BT, D’Esposito M (2005) Searching for “the top” in top-down control. Neuron 48:535–538.

Miller EK, Cohen JD (2001) An integrative theory of prefrontal cortex function. Annual Review of Neuroscience 24:167–202.

Moore CI, Carlen M, Knoblich U, Cardin JA (2010) Neocortical interneurons: From diversity, strength. Cell 142:189–193.

Nelissen N, Stokes M, Nobre AC, Rushworth MF (2013) Frontal and parietal cortical interactions with distributed visual representations during selective attention and action selection. J Neurosci 33:16443–16458.

Norman-Haignere SV, McCarthy G, Chun MM, Turk-Browne >NB (2012) Category-selective background connectivity in ventral visual cortex. Cereb Cortex 22:391–402.

O’Craven KM, Downing PE, Kanwisher N (1999) Fmri evidence for objects as the units of attentional selection. Nature 401:584–587.

Power JD, Barnes KA, Snyder AZ, Schlaggar BL, Petersen SE (2012) Spurious but systematic correlations in functional connectivity mri networks arise from subject motion. NeuroImage 59:2142–2154.

Ruff CC, Bestmann S, Blankenburg F, Bjoertomt O, Josephs O, Weiskopf N, Deichmann R, Driver J (2008) Distinct causal influences of parietal versus frontal areas on human visual cortex: Evidence from concurrent tms-fmri. Cereb Cortex 18:817–827.

Saalmann YB, Pinsk MA, Wang L, Li X, Kastner S (2012) The pulvinar regulates information transmission between cortical areas based on attention demands. Science 337:753–756.

Seidl KN, Peelen MV, Kastner S (2012) Neural evidence for distracter suppression during visual search in real-world scenes. J Neurosci 32:11812–11819.

Serences JT, Saproo S, Scolari M, Ho T, Muftuler LT (2009) Estimating the influence of attention on population codes in human visual cortex using voxel-based tuning functions. Neuroimage 44:223–231.

Sherman SM (2016) Thalamus plays a central role in ongoing cortical functioning. Nat Neurosci 16:533–541.

Shine JM, Koyejo O, Poldrack RA (2016a) Temporal metastates are associated with differential patterns of time-resolved connectivity, network topology, and attention. Proc Natl Acad Sci U S A 113:9888–9891.

Shine JM, Koyejo O, Bell PT, Gorgolewski KJ, Gilat M, Poldrack RA (2015) Estimation of dynamic functional connectivity using Multiplication of Temporal Derivatives. Neuroimage 122:399–407.

Shine JM, Bissett PG, Bell PT, Koyejo O, Balsters JH, Gorgolewski KJ, Moodie CA, Poldrack RA (2016b) The Dynamics of Functional Brain Networks: Integrated Network States during Cognitive Task Performance. Neuron 92:544–554.

Smith SM, Jenkinson M, Woolrich MW, Beckmann CF, Behrens TE, Johansen-Berg H, Bannister PR, De Luca M, Drobnjak I, Flitney DE, Niazy RK, Saunders J, Vickers J, Zhang Y, De Stefano N, Brady JM, Matthews PM (2004) Advances in functional and structural mr image analysis and implementation as fsl. NeuroImage 23 Suppl 1:S208–219.

Treue S, Trujillo >JCM (1999) Feature-based attention influences motion processing gain in macaque visual cortex. Nature 399:575–579.

Van Essen DC, Anderson CH, Felleman DJ (1992) Information processing in the primate visual system: An integrated systems perspective. Science 255:419–422.

Vierling-Claassen D, Cardin JA, Moore CI, Jones SR (2010) Computational modeling of distinct neocortical oscillations driven by cell-type selective optogenetic drive: Separable resonant circuits controlled by low-threshold spiking and fast-spiking interneurons. Front Hum Neurosci 4:198.

Wang X-J (2010) Neurophysiological and computational principles of cortical rhythms in cognition. Physiol Rev 90:1195–1268.

Willenbockel V, Sadr J, Fiset D, Horne GO, Gosselin F, Tanaka JW (2010) Controlling low-level image properties: the SHINE toolbox. Behav Res Methods 42:671–684.

Wimmer RD, Schmitt LI, Davidson TJ, Nakajima M, Deisseroth K, Halassa MM (2015) Thalamic control of sensory selection in divided attention. Nature 526:705–709.

Zanto TP, Rubens MT, Thangavel A, Gazzaley A (2011) Causal role of the prefrontal cortex in top-down modulation of visual processing and working memory. Nat Neurosci 14:656–661.

